# The *Botrytis cinerea* Gene Expression Browser

**DOI:** 10.1101/2022.08.29.504976

**Authors:** Gabriel Pérez-Lara, Tomás C. Moyano, Andrea Vega, Luis F. Larrondo, Rubén Polanco, José M. Álvarez, Daniel Aguayo, Paulo Canessa

## Abstract

To analyze and visualize comprehensive gene expression patterns in the phytopathogenic fungus *Botrytis cinerea*, we developed BEB — a web-based *B. cinerea* gene expression browser. This tool and associated databases (DB) contain manually-curated RNA-Seq experiments conducted in *B. cinerea*. BEB allows easy gene expression analyses of genes of interest under different culture conditions by providing publication-ready heatmaps depicting transcripts levels. BEB is a computationally-inexpensive web-based application and gene expression DB that allows effortless visualization of the transcript levels of genes of interest without needing advanced computational skills. BEB also provides details of each experiment under analysis and user-defined gene expression clustering and visualization options. If needed, tables of gene expression values can be downloaded for further exploration, employing more sophisticated bioinformatics tools. The BEB implementation is based on open-source computational technologies that can be easily deployed for other organisms of interest with little additional effort. To demonstrate BEB’s usability and potential, we selected genes of interest in *B. cinerea* to determine their expression patterns across different conditions. We thus focused our analysis on secondary metabolite gene clusters, chromosome-wide gene expression, previously described virulence factors, and reference genes, leading to a comprehensive expression overview of these groups of genes in this relevant fungal phytopathogen.

## Introduction

Genome-scale collection of gene expression data in RNA sequencing (RNA-Seq) is refashioning modern molecular biology strategies. Biological models across phyla have benefited from the most recent technological advances in today’s various massive sequencing methodologies including short and emerging long-read transcriptomics (Stark et al., 2019). This can facilitate accurate gene expression profiling in virtually any organism and experimental condition. RNA-Seq thus represents a tool that can provide important clues regarding the function and regulation of different genes of interest (Kukurba and Montgomery, 2015).

The standard workflow of RNA-Seq experiments relies on high-quality RNA extraction. After adequate quantity and quality assessment of the nucleic acid under analysis (Sheng et al., 2016), researchers build sequencing libraries following rigorous and standardized methods that ensure the most out of each sequence read. Nonetheless, low-quality reads and adapter sequences must be discarded before differential gene expression analyses (Bolger et al., 2014). These procedures allow downstream time-consuming transcriptome mapping of each sequence to a reference genome. Researchers thus employ specialized short read alignments tools such as STAR (Dobin et al., 2012), TopHat2 (Kim et al., 2013), Hisat2 (Kim et al., 2015), or Kallisto (Bray et al., 2016), among others, to quantify each read (Liao et al., 2019; Anders et al., 2014) to ultimately apply a suitable differential expression detection algorithm (Love et al., 2014; Chen et al., 2016). Readers can consult several reviews to address many critical considerations at each step (Stark et al., 2019; Dorado et al., 2021; Hrdlickova et al., 2017; Bayega et al., 2018). Ironically, after all these massive scientific and computational efforts to make RNA-Seq data biologically accurate, meaningful, and accessible to most biologists, RAW sequence files are deposited back into public databases such as NCBI’s Sequence Read Archive (SRA) (Leinonen et al., 2011). Therefore, there is a significant scientific, technical, and computational challenge when scientists with no bioinformatics expertise nor computational power seek to analyze all across-laboratory experiments to determine otherwise hidden global gene expression patterns.

Several initiatives — most of them in well-known model species — have been propelled to circumvent some of the abovementioned difficulties. These include the model plant species *Arabidopsis thaliana* (Sullivan et al., 2019), agricultural relevant plants (Robinson et al., 2018), and several other species aggregated in massive initiatives like the “Expression Atlas” of the European Bioinformatics Institute (EMBL-EBI) (Papatheodorou et al., 2019). This latter collection holds information on 22 animal models, over nine plant species different from *Arabidopsis*, and only three fungal species, with 95.7% of the RNA-Seq experiments concentrated in the budding yeast *Saccharomyces cerevisiae*. Therefore, there is a lack of implementation of this kind of tool and curated gene expression information in fungal species. Undoubtedly, this represents an opportunity to better understand the biology of this relevant but often neglected group of organisms (Case et al., 2020).

One notable exception is the wheat fungal pathogen *Puccinia striiformis* f. sp. *tritici*, with its recently developed platform for analyzing gene expression patterns in a myriad of culture conditions including “*in planta*” growth (Adams et al., 2021). This strategy can provide meaningful insights regarding, for instance, the infection strategies employed by this pathogen. While fungi represent an exceptional biotechnological chassis, their extraordinary adaptation capacity to diverse environmental niches also means several risks for animal health and agricultural production (Case et al., 2020; Fisher et al., 2020). In fact, ten fungal phytopathogens have long been considered highly relevant agricultural threats. Unfortunately, despite the availability of a relatively small but significant number of transcriptomics experiments for most of them **(Table 1)**, there is no simple and easy-to-use tool to determine gene expression patterns. With a great degree of host specificity (Couch et al., 2005), the most important specialist phytopathogenic fungus is *Magnaporthe oryzae*: the causal agent of the rice blast disease. On the other hand, the most relevant generalist is the so-called grey mold fungus *Botrytis cinerea* (Dean et al., 2012). Both have an enormous negative impact on food security and production worldwide.

**Table 1.** Publicly available RNA-Seq experiments and studies in the NCBI’s short read archive (SRA) for the top 10 most relevant fungal phytopathogens (Dean et al., 2012).

| Fungal Species | SRA Experiments | SRA Studies |
| --- | --- | --- |
| *Magnaporthe oryzae* | 1714 | 125 |
| *Botrytis cinerea* | 2403 | 89 |
| *Puccinia spp* | 3847 | 195 |
| *Fusarium graminearum* | 2141 | 177 |
| *Fusarium oxysporum* | 2700 | 188 |
| *Blumeria graminis* | 1057 | 43 |
| *Mycosphaerella graminicola* | 2095 | 230 |
| *Colletotrichum spp* | 1752 | 219 |
| *Ustilago maydis* | 538 | 38 |
| *Melampsora lini* | 205 | 3 |

*B. cinerea* is the most largely investigated necrotrophic fungal plant pathogen. It has been the focus of numerous research groups over several decades. This single fungal species explains over $10 billion in agricultural product losses (Weiberg et al., 2013). In the *Botrytis* genus, several species are specialist plant pathogens (Staats et al., 2005). In contrast, with its necrotrophic infection strategy, *B. cinerea* can infect over 1000 plant species (Veloso and van Kan, 2018). For the interested reader, there are several seminal works revisiting canonical *B. cinerea* infection strategies (van Kan, 2006; Choquer et al., 2007; Shlezinger et al., 2011); contemporary research trends and most current advances have been reviewed elsewhere (Mbengue et al., 2016; Castillo et al., 2017; Schumacher, 2017; Veloso and van Kan, 2018; Larrondo and Canessa, 2019; Cheung et al., 2020).

Since the foundational analyses that facilitated the first genome database of *B. cinerea* (Amselem et al., 2011), several improvements have been conducted over the years (Staats and van Kan, 2012). This led to a gapless genome whose assembly was supported with an optical map (van Kan et al., 2017). These genomic advances have allowed the accumulation of a significant number of transcriptomic experiments **(Table 1)** that largely remain underexplored due to the lack of tools to analyze all expression data simultaneously.

To visualize organism-wide gene expression patterns in *B. cinerea*, we developed the *B. cinerea* gene Expression Browser (BEB). With a user-friendly interface, this tool allows straightforward gene expression analysis of genes of interest under various conditions. For this purpose, users only need to provide *B. cinerea* gene IDs. To demonstrate the usability and potential of this tool, we picked several genes of interest, including virulence factors, to determine their expression patterns across experimental conditions.

## Materials and methods

### RNA-Seq datasets available for *Botrytis cinerea* global gene expression analysis

To generate a robust web-based platform capable of visualizing global gene expression patterns of *B. cinerea* across available genome-wide expression experiments, we put together all publicly RNA-Seq data retrieved from the NCBI’s Sequence Read Archive (SRA) database as well as the EMBL-EBI (European Bioinformatics Institute) available on November 30, 2021. The uploaded dataset was composed of 218 individual files representing 76 experimental groups. RNA-Seq experiments included, but are not limited to, those with *B. cinerea* growing in axenic *in vitro* cultures (non-infective conditions; i.e., *on plate* and liquid medium) and during the infection of different plant species (dual RNA-Seq; i.e., *B. cinerea* infecting *A. thaliana*, among others). Details are provided in **Supplementary Material 1**.

### Gene expression metadata construction

The manually-curated metadata available on the BEB **(Supplementary Material 1)** uses the NCBI SRA’s metadata information schema (Leinonen et al., 2010) and describes the general experimental conditions of each RNA-Seq experiment including replicates (described as group_for_averaging), available treatments (description of the culture conditions in which the experiment was performed), utilized *B. cinerea* strain/isolate, the type of tissue, and, when appropriate, the presence and type of plant material infected by the fungus (e.g., tissue and hours post-infection, when available). Importantly, not all RNA-Seq available in NCBI SRA’s contained a complete description of the experimental condition (e.g., information to cross-check the data with the associated sequencing file). Thus, when possible, we also manually analyzed all associated publications making every effort to obtain as much information as possible. If available, PubMed IDs of the respective publication were also included. Those RNA-Seq experiments that were impossible to determine the FASTQ file confidently and its respective experimental condition were not included. Studies focused on small RNA were not included. Importantly, the format used in **Supplementary Material 1** is consistent with the CSV schema used by BEB’s server (see below).

### Data pre-processing and RNA-Seq experiments mapping

As RNA-Seq data was composed of both Single (SE) and Paired-End (PE) Illumina sequencing technologies, a careful examination of the data was performed before mapping. First, a quality inspection was assessed employing fastQC (version 0.11.8, (Wingett and Andrews, 2018)). After this procedure, low-quality reads and sequencing adapters from each FASTQ files were filtered out using BBDuk (https://sourceforge.net/projects/bbmap/) (v38.90; ktrim=r k=23 mink=11 hdist=2 qtrim=rl trimq=10 ftm=5 maq=15 minlength=50 tbo). Thereafter, filtered reads were pseudoaligned to the *B. cinerea* B05.10 transcriptome (van Kan et al., 2017) (ASM83294v1) using Kallisto (v0.46.0) (Bray et al., 2016). Kallisto SE mapping was performed under the following settings: –single -b 100 -l 100 -s 20. Standard parameters (-b 100) were employed for PE. The *B. cinerea* transcriptome reference was downloaded from EnsemblFungi release 52 (Howe et al., 2019) representing the previously published work (van Kan et al., 2017).

### Gene expression analysis and BEB’s transcriptional profile experiments database

Kallisto’s mapped read counts were further processed to infer transcript abundances with the tximport package in R (Soneson et al., 2015) (v1.20.0; on RStudio v4.1.0). This approach enabled us to obtain a complete dataset containing gene-level estimated counts derived from all RNA-Seq experiments. A custom Python version of the DESeq2 median of the ratio normalization method was used to determine gene expression levels (Anders and Huber, 2010; Love et al., 2014). The gene-level count matrix and the metadata file described above are loaded and available to explore on BEB’s server database and browser (see below).

### BEB’s server implementation and user interface

The BEB’s server is a web-based tool written in Python 3.7 using Streamlit’s open-source app framework and Docker. The BEB’s metadata and code were used to prepare the data, set up, and run the web-based tool presented herein are available on GitHub (https://github.com/ibioChile/CanessaLab). A working version is available at https://beb.canessalab.org. The BEB’s landing page contains a left sidebar section where the experimental factors — extracted from the experimental metadata file — can be selected through dropdown lists. In addition, a list of genes of interest can be used as an entry on BEB’s landing page or these can be randomly selected from BEB’s dataset **(Figure 1)**. Importantly, gene identifiers must be provided separated by spaces (in the form of Bcin[XX]g[YYYYY]; where “XX” corresponds to the chromosome and “YYYYY” to the gene number). Once parameters are selected and submitted, the bottom section shows a customizable colored heatmap. This heatmap depicts the expression levels of the provided subset of genes in the experiments that fulfill the selected factors. The heatmap customization parameters include coloration of the expression levels by quartiles, DESeq2 units, or log2 transformation to highlight fold differences among experiments. Furthermore, both genes and experiments can be clustered to help the user identify co-expressed gene and expression trends.

**Figure 1.**
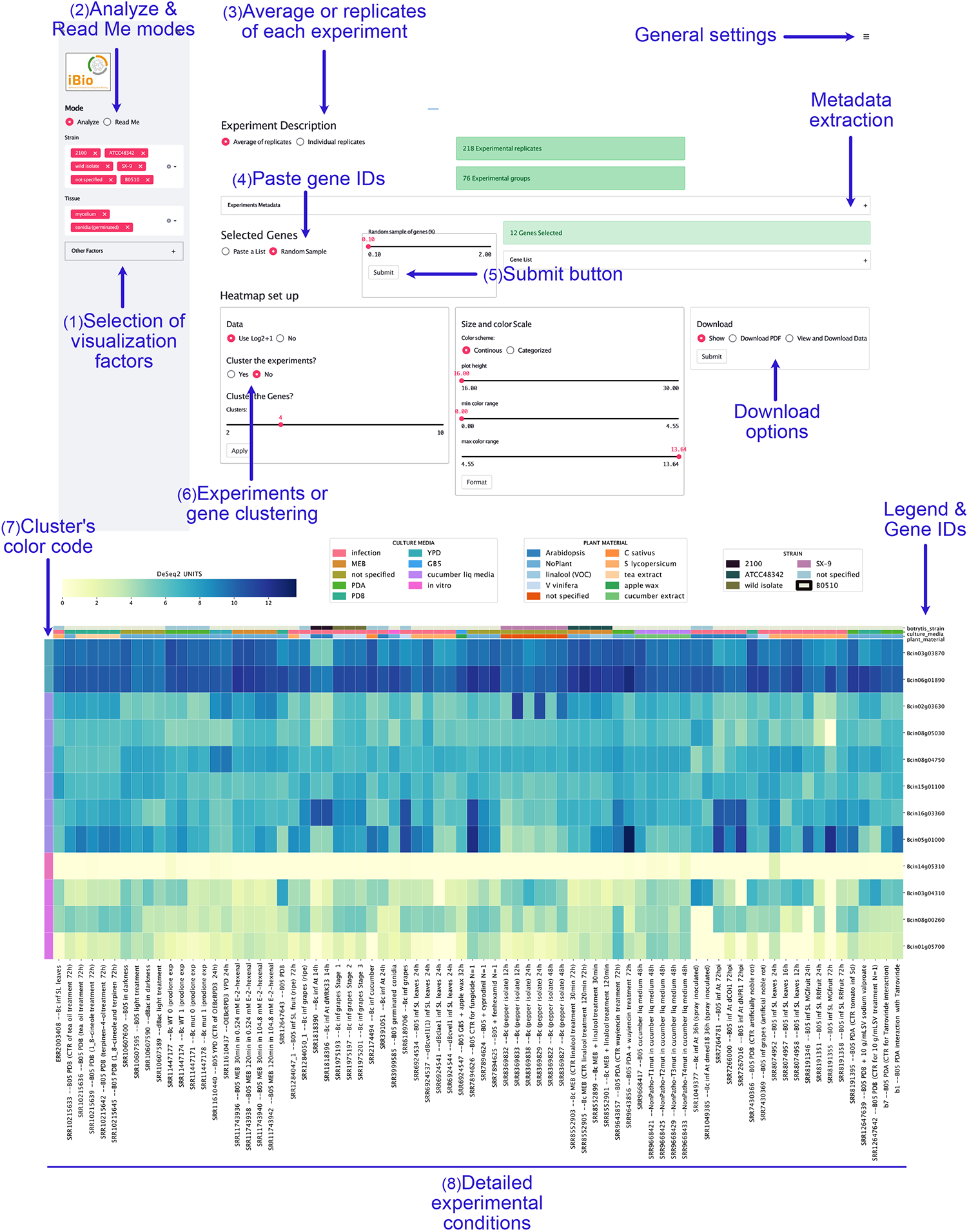
*B. cinerea* gene expression browser (BEB) graphical user interface. The BEB’s landing page contains a left sidebar section where experimental factors can be selected through dropdown lists (1). The “Analyze” and “Read Me” options (2) are available on the left sidebar (top section). Detailed instructions on how to use BEB are available in the latter display tool. In the upper section, the user can select to either display average expression values or each replicate individually (3). Users can copy, paste, and submit gene IDs in the right middle section after selecting the “Paste a List” option (4). After clicking the “Submit” button (5), a heatmap depicting gene expression is generated at the bottom of the webpage. Clustering and download options are available in the right middle section (6). BEB’s clusters are denoted with a color code located at the outmost left column of the heatmap (7). A detailed description of each experiment is provided at the bottom of the figure including SRA IDs for reference (8).

### Additional bioinformatic analyses

To predict secondary metabolite gene clusters in *B. cinerea*, antiSMASH (version 6.1.1) (Blin et al., 2019) software was used employing default parameters. Succinctly, FASTA and GFF3 files from the *B. cinerea*’s genome database were provided, and the output was manually inspected. For the genes encoded in chromosomes 17 and 18, we also performed a BLAST2GO (Götz et al., 2008) automatic analysis to retrieve all available functional annotations.

Due to the quantity of expression data deposited in BEB, we also looked for putative/new reference genes that can be used for future reverse transcription quantitative real-time PCR (RT-qPCR) studies. For this purpose, read counts of each *B. cinerea* gene were normalized with the total mapped reads per library. An additional normalization was then performed by the upper quartile and median norm, as described for RNA-Seq data (Carmona et al., 2017). Finally, the normalized reads of each gene were standardized by the transcript size and classified by quartiles of coefficient of variation. Likely reference genes have the lowest coefficient of variation as demonstrated previously (Carmona et al., 2017; Pombo et al., 2017; Tilli et al., 2016).

## Results and discussion

### Global gene expression patterns of phytotoxic secondary metabolite gene clusters in *B. cinerea*

Biosynthetic gene clusters (BGCs) in *B. cinerea* are common. Since the first version of its genome project (Amselem et al., 2011), at least 40 groups of genes that orchestrate the synthesis of secondary metabolites (SM) have been identified. One of these SM is botcinic acid, a phytotoxic polyketide produced by the coordinated action of a gene cluster in a subtelomeric region of chromosome (Chr) 1 (Dalmais et al., 2011). The transcription factor (TF) BcBoa13 **(Table 2)** controls this cluster’s transcriptional regulation (Porquier et al., 2019). The BEB-generated heatmap plot for these genes **(Table 2, Figure 2A)** shows low expression in PDA, PDB (Potato Dextrose Agar or Broth, respectively), YPD (Yeast Extract–Peptone–Dextrose), and MEB (malt extract broth) (*in vitro*) culture media. The highest values were seen during the infection of plants including *Solanum lycopersicum* and *A. thaliana* (middle section of Figure 2A). This observation is consistent with previous studies showing the induction of botcinic acid’s genes in the process of infection (Porquier et al., 2019). The most notable exceptions were gene IDs Bcin01g00150, Bcin01g00160, and Bcin01g00170 (depicted at the top of **Figure 2A**). These genes are physically located at one of the cluster’s borders.

**Table 2.** Gene IDs and names of the two major phytotoxic secondary metabolites of *B. cinerea*. Functional descriptions derived from manual inspection of the *B. cinerea* genome database and/or their respective publication.

| Gene ID | Name | Functional description (manual and/or from publications) |
| --- | --- | --- |
| Bcin01g00010 | Bcboa1 | Putative NmrA-like regulator |
| Bcin01g00020 | Bcboa2 | Putative flavin-binding monooxygenase-like |
| Bcin01g00030 | Bcboa3 | Putative monooxygenase |
| Bcin01g00040 | Bcboa4 | Putative monooxygenase |
| Bcin01g00050 | Bcboa5 | Putative alcohol dehydrogenase |
| Bcin01g00060 | Bcboa6,Bcpks6 | Polyketide synthase (Dalmais et al., 2011); Secondary Metabolism Key Enzyme |
| Bcin01g00070 | Bcboa7 | Putative monooxygenase |
| Bcin01g00080 | Bcboa8 | Putative FAD-binding protein |
| Bcin01g00090 | Bcboa9,Bcpks9 | Polyketide synthase (Dalmais et al., 2011); Secondary Metabolism Key Enzyme |
| Bcin01g00100 | Bcboa10 | Putative thioesterase |
| Bcin01g00110 | Bcboa11 | Putative transferase |
| Bcin01g00120 | Bcboa12 | Unknown |
| Bcin01g00130 | Bcboa13 | Zn(II)2Cys6 transcription factor (Porquier et al., 2019) |
| Bcin01g00140 | Bcboa15 | Pseudogene (Porquier et al. 2019) |
| Bcin01g00150 | Bcboa16 | Putative dehydratase/ Pseudogene (Porquier et al., 2019) |
| Bcin01g00160 | Bcboa17 | Putative dehydrogenase |
| Bcin01g00170 | no name | Putative permease (Porquier et al., 2019) |
| Bcin12g06370 | Bcbot4 | Cytochrome P450 monooxygenase |
| Bcin12g06380 | Bcbot1 | Benzoate 4-monooxygenase cytochrome p450 |
| Bcin12g06390 | Bcbot2,Bcstc1 | Sesquiterpen cyclase |
| Bcin12g06400 | Bcbot3 | Cytochrome P450 monooxygenase |
| Bcin12g06410 | Bcbot5 | Acetyl transferase, in botrydial biosynthesis gene cluster |
| Bcin12g06420 | Bcbot6 | Zn(2)-C6 fungal-type transcription factor, (Porquier et. al., 2016) |

**Figure 2.**
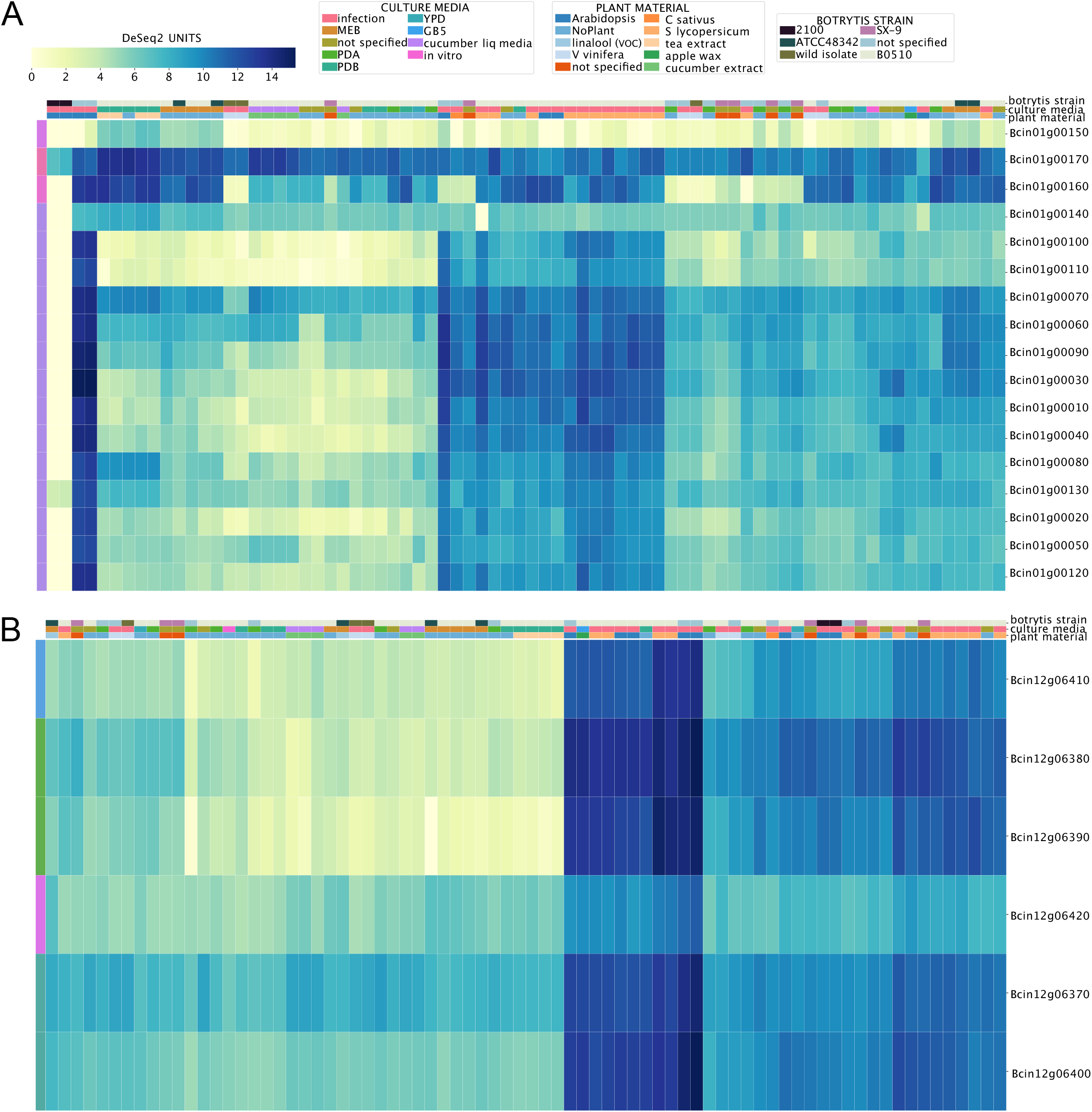
Expression patterns of the gene clusters involved in the production of botcinic (A) and botrydial (B) acid phytotoxins in *B. cinerea*. The heatmaps depict mRNA levels as calculated employing DeSeq2 (see Methods). A color scale distinguishes each gene expression level as low or highly expressed (from yellow to dark blue, respectively). A color code at the top of each heatmap is also used to indicate culture media, plant material, and *B. cinerea* strains (legend at the top). Experimental conditions are indicated in each column, while analyzed genes are in each row. To streamline the overall figure, the description of each experiment was omitted in both heatmaps (compared with Figure 1). BEB’s clusters are denoted with a color code located at the outmost left column of each heatmap.

Another relevant BGC in *B. cinerea* is required for botrydial production, an additional phytotoxic SM synthesized by this fungus. When explored on the BEB, the *bot* genes needed for botrydial synthesis show a similar expression pattern as the one observed for botcinic acid, with higher mRNA levels during the infection of plant tissue (see **Figure 2B)**. BEB’s gene clustering algorithm (see methods) is denoted with a color code at the outmost left column of the respective heatmap and facilitates the recognition of distinctive gene expression patterns under different conditions. It also allows for capturing distinctive patterns within genes. For example, the TF BcBOT6 (Bcin12g06420), which is central for the biosynthesis of botrydial (Porquier et al., 2016), displays the most distinctive expression pattern compared with the five non-regulatory clusters’ genes.

### Gene expression of orphan secondary metabolite gene clusters

While several genes encoding enzymes required for SM synthesis have been identified in the genome of *B. cinerea* **(Table 3)**, a myriad of them are predicted to participate in synthesizing unknown compounds (Sabine Fillinger, 2016). Since the expression pattern of these genes is unknown, we decided to use the BEB to analyze them and determine whether this tool can shed light on experimental conditions that could facilitate the study of SM biosynthesis. Among sesquiterpene cyclase encoding genes, Bcin12g06390 (*bcbot2*; botrydial, see above; **Table 3**) displayed the highest expression values during the infection of *A. thaliana* and *S. lycopersicum*, which is clearly visualized using the BEB’s quartile-categorized expression option **(Figure 3A)**. For comparative purposes, the continuous (color) scale is also displayed (**Figure 3B)**. Most of the remaining genes showed relatively low expression values with Bcin01g03520 and Bcin04g03550 the most notable exceptions. Interestingly, the latter gene appears highly expressed during the infection of *S. lycopersicum* and *in vitro* cultures supplemented with cucumber or tea extracts.

**Table 3.** Gene IDs and names of key enzymes required for the synthesis of secondary metabolites (SM) in *B. cinerea* as described previously (Sabine Fillinger, 2016). The associated SM were inferred from each gene putative function or their respective publication.

| Gene ID | KE name(s) | Metabolite |
| --- | --- | --- |
| Bcin12g06390 | BcStc1/Bot2 | Botrydial (Pinedo et al. 2008) |
| Bcin08g02350 | BcStc2 | Unknown sesquiterpene |
| Bcin13g05830 | BcStc3 | Unknown sesquiterpene |
| Bcin04g03550 | BcStc4 | Unknown sesquiterpene |
| Bcin01g03520 | BcStc5/BcAba5 | Unknown sesquiterpene |
| Bcin01g04920 | BcDtc1 | Unknown diterpene |
| Bcin08g03560 | BcDtc3 | Unknown diterpene |
| Bcin05g05670 | BcPax1 | Unknown indole-diterpene |
| Bcin01g04560 | BcPhs1 | Retinal (Schumacher et al. 2014), phytoene synthase |
| Bcin01g03260 | BcBik1 | Bikaverin (Schumacher et al. 2013) |
| Bcin01g00060 | BcPks6/Boa6 | Botcinic acid (Dalmais et al. 2011) |
| Bcin01g00090 | BcPks9/Boa9 | Botcinic acid (Dalmais et al. 2011) |
| Bcin02g08770 | BcPks12 | Melanin (Schumacher et al. 2014) |
| Bcin03g08050 | BcPks13 | Melanin (Schumacher et al. 2014) |
| Bcin14g00600 | BcPks1 | Unknown polyketide |
| Bcin02g01680 | BcPks2 | Unknown polyketide |
| Bcin11g02700 | BcPks4 | Unknown polyketide |
| Bcin07g02920 | BcPks8 | Unknown polyketide |
| Bcin13g01510 | BcPks10 | Unknown polyketide |
| Bcin14g01290 | BcPks11 | Unknown polyketide |
| Bcin16g01830 | BcPks14 | Unknown polyketide |
| Bcin05g06220 | BcPks15 | Unknown polyketide |
| Bcin16g05040 | BcPks16 | Unknown polyketide |
| Bcin03g02010 | BcPks17 | Unknown polyketide |
| Bcin02g08830 | BcPks18 | Unknown polyketide |
| Bcin08g00290 | BcPks19 | Unknown polyketide |
| Bcin04g00640 | BcPks20 | Unknown polyketide |
| Bcin05g08400 | BcPks21 | Unknown polyketide |
| Bcin13g02130 | BcChs1/Bpks | Pyrones, resorcylic acids and resorcinols (Jeya et al. 2012) |
| Bcin12g00690 | BcNrps2 | Ferrichrome siderophores (Bushley and Turgeon 2010) |
| Bcin16g03570 | BcNrps3 | Ferrichrome siderophores (Bushley and Turgeon 2010) |
| Bcin01g11450 | BcNrps7 | Ferrichrome siderophores (Bushley and Turgeon 2010) |
| Bcin01g03730 | BcNrps6 | Coprogene siderophore (Bushley and Turgeon 2010) |
| Bcin12g04980 | BcNrps1 | Unknown peptides |
| Bcin02g02380 | BcNrps4 | Unknown peptides |
| Bcin04g01390 | BcNrps5 | Unknown peptides |
| Bcin11g02650 | BcNrps8 | Unknown peptides |
| Bcin14g01300 | BcNrps9 | Unknown peptides |
| Bcin03g04360 | BcPks3 | Unknown amino-acid containing- polyketides (PKS-NRPS hybrids) |
| Bcin01g11550 | BcPks5 | Unknown amino-acid containing- polyketides (PKS-NRPS hybrids) |
| Bcin10g00040 | BcPks7 | Unknown amino-acid containing- polyketides (PKS-NRPS hybrids) |
| Bcin16g01940 | Bcdmats1 | Unknown alkaloids |

**Figure 3.**
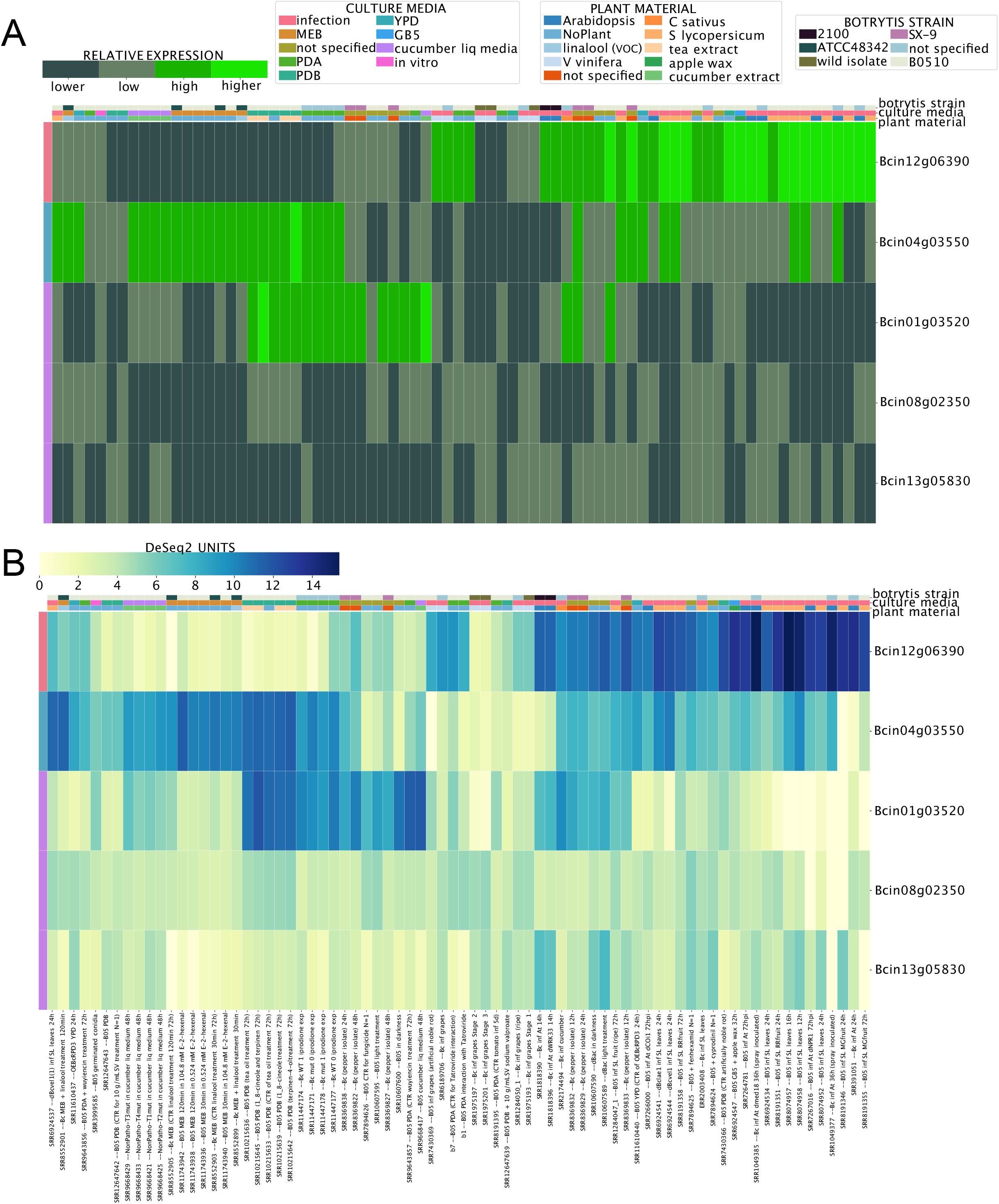
Expression patterns of five sesquiterpene cyclase enzyme encoding genes. **(A)** Heatmap depicting the mRNA expression levels employing the BEB’s quartile-categorized expression option (lower: dark green; higher: light green). The continuous color scale **(B)** from low to high expression (yellow to dark blue, respectively) is also shown for comparative purposes. A color code at the top denotes culture conditions and *B. cinerea* strains. Experimental conditions and analyzed genes are indicated in columns and rows, respectively. Gene IDs are indicated at the right of each heatmap. A detailed description of each experiment (for both heatmaps) is provided at the bottom of the figure including SRA IDs for reference. Gene expression clusters are denoted with a color code located at the outmost left column of each heatmap.

Among the polyketide synthases (PKS, **Table 3**), Bcin01g00060 and Bcin01g00090 (required for botcinic acid biosynthesis, see above) display the highest expression values during the infection of *A. thaliana* and *S. lycopersicum*, as shown in **Figure 4A**. In contrast, seven PKS genes (shown in the middle-bottom left of **Figure 4A**) display low expression values in most culture conditions, with particularly low mRNA levels during the infection of tomato plants. Interestingly, among diterpene cyclases **(Figure 4B, Table 3)**, Bcin01g04920 showed the highest expression values during the infection of cucumber plants or liquid media supplemented with cucumber extract (indicated by asterisks in **Figure 4B**). These observations show how BEB can be used to generate new hypotheses, i.e., an experimental condition that could be used to study the function of this particular gene.

**Figure 4.**
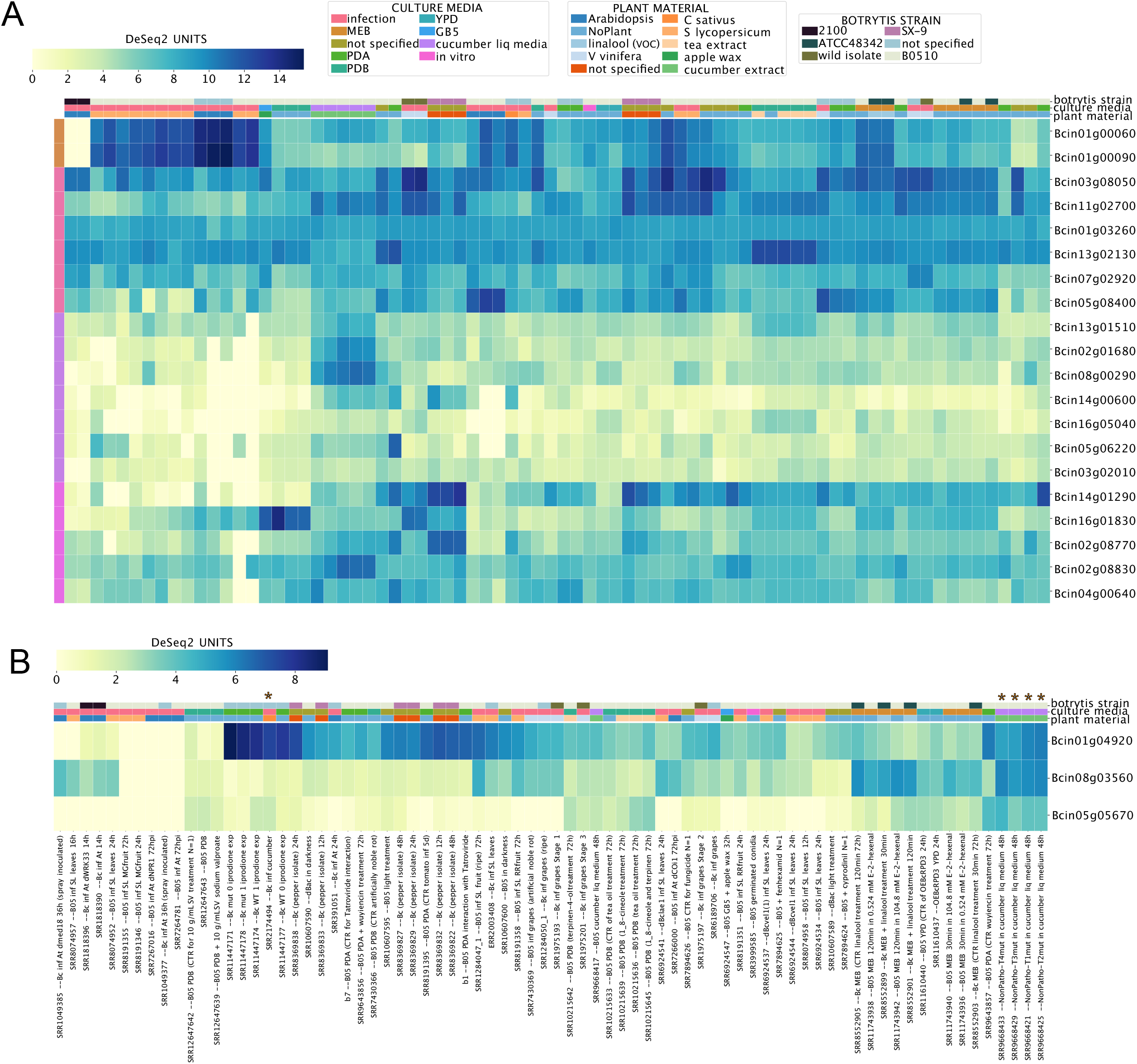
Polyketide synthases (A, PKS) and diterpene cyclases (B) expression levels across BEB experimental conditions. Both heatmaps indicate transcript expression levels with their corresponding expression scale from yellow to dark blue (low or high expression, respectively). The legend at the top of the figure denotes culture conditions and *B. cinerea* strains. Experimental conditions and genes are depicted in columns and rows, respectively. Gene IDs are provided at the right of each heatmap. Gene expression clusters are denoted with a color code located at the outmost left column of each heatmap. For (B), a detailed description of each experiment is provided at the bottom of the figure. Columns (in B) denoted with asterisks are discussed in the main text.

Finally, we looked at the expression values of NRPS (non-ribosomal peptide synthetases) and hybrid PKS-NRPS encoding genes **(Table 3)**. Gene IDs Bcin01g11450 (putatively involved in ferrichrome siderophores biosynthesis) and Bcin01g11550 (unknown polyketide) showed a very similar expression pattern among PKS-NRPS genes, as indicated by BEB’s clustering function (outmost left colored column of **Figure 5A**). These two genes were clustered in a genomic region spanning *circa* 45kbp depicted in **Figures 5B and 5C** (Region 5), a subtelomeric region in Chr 1 opposite to the botcinic acid SM cluster (**Figure 5B**, Region 1). The physical proximity of gene IDs Bcin01g11450 and Bcin01g11550 and their particular expression pattern observed through BEB’s heatmaps encouraged us to investigate whether all genes within the 45kbp region may represent a putative BGC. According to the antiSMASH software (Blin et al., 2019) that allows *in silico* identification of SM gene clusters in fungal systems, the region mentioned above corresponds to one out of 5 SM clusters in *B. cinerea*’s Chr 1, predicted between genomic coordinates 4,002,878-4,093,700 **(Figure 5B)**. To support this prediction, we analyzed the expression pattern of all genes **(Figure 5C)** encoded in this putative BGC using BEB. All genes display highly similar expression patterns across all culture conditions **(Figure 5D)**. This suggests co-regulation as expected from an SM gene cluster, most likely involved in ferrichrome siderophore biosynthesis. Since BGCs usually contain a transcription factor (TF) encoded within these collections of genes, we analyzed if such a regulatory protein might be present in Region 5 **(Figure 5C)**. Indeed, employing a manually-curated catalog of TF for *B. cinerea* (Olivares-Yañez et al., 2021), along with careful examination led to the Bcin01g11510 gene encoding a fungal Zn(2)-Cys(6) binuclear TF whose participation in the regulation of this BGC or siderophore biosynthesis has not been previously described. Interestingly, this TF gene displays the most distinctive expression pattern among all genes in the SM biosynthesis cluster as indicated by the BEB’s gene clustering function (outmost left colored column of **Figure 5D**). Siderophores are complex low-molecular-weight molecules involved in iron acquisition. Little is known about *B. cinerea*’s iron acquisition pathways and transcriptional regulatory mechanisms. However, this fungus is expected to synthesize at least nine siderophores to support this metal acquisition (Konetschny-Rapp et al., 1998; Bushley and Turgeon, 2010) in addition to the membrane-bound reductive iron assimilation mechanism that also participates in iron incorporation (Vasquez-Montaño et al., 2020).

**Figure 5.**
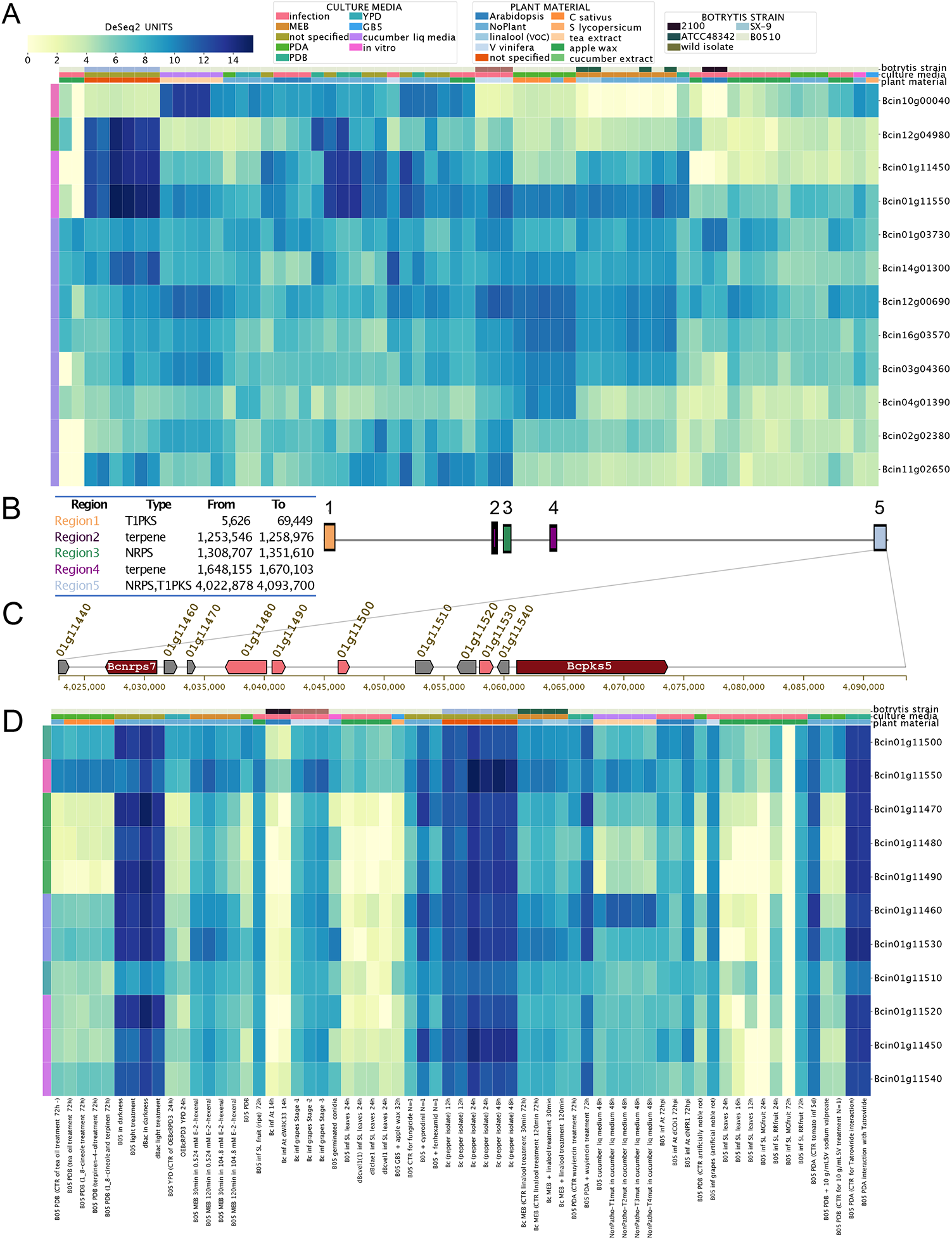
Gene expression patterns of non-ribosomal peptide synthetases (NRPS) and hybrid polyketide synthases (PKS-NRPS) encoding genes in *B. cinerea*. **(A)** Expression patterns of all predicted NRPS and PKS-NRPS mentioned in Table 3. **(B)** Identification of the five SM gene clusters in *B. cinerea*’s chromosome (Chr) 1. The table inset describes each of the five SM regions within Chr1, the predicted SM type, and their respective genomic coordinates. Colored boxes represent each region (right, with numbers). **(C)** Clustered genes within “Region 5” as described in (B). Boxes with arrowheads indicate each gene’s transcriptional orientation within the SM gene cluster. Dark red and pink arrowhead boxes indicate core and additional biosynthetic genes, respectively. Grey indicates other genes. Gene IDs and gene names are indicated. For simplicity, the “Bcin” prefix was omitted. **(D)** Expression patterns of all genes encoded in “Region 5” as indicated in (B) and (C), from *bcnrps7* to *bcpks5*. Both heatmaps indicate mRNA levels and experimental conditions as described in the former figures. In (D), a detailed description of each experiment is provided at the bottom of the heatmap.

These findings collectively illustrate the BEB’s ability to reveal the expression patterns of orphan gene clusters. This information can facilitate their investigation under specific experimental conditions. Combined with easy-to-use tools such as antiSMASH (Blin et al., 2019) and up-to-date TF databases (Olivares-Yañez et al., 2021), the gene expression patterns determined in BEB could also provide testable hypotheses regarding transcriptional regulation.

### Chromosome-wide gene expression analysis

The latest iteration of the genome sequencing process of the *B. cinerea* B05.10 strain revealed exciting structural features including two previously unnoticed minichromosomes (van Kan et al., 2017). In *B. cinerea*, this particular group of Chr 17 and 18 comprises 18 and 16 protein-encoding genes, respectively. This is a common feature observed in these genetic arrangements. These accessory chromosomes (AC) are generally small and are not considered essential for the organisms’ survival. Among some other pathogenic fungi, they are characterized by the presence of genes encoding virulence factors (Ma et al., 2010; van Dam et al., 2017; Li et al., 2020). The *B. cinerea* genes encoded in ACs display little or no similarity to other genes in other organisms including fungi (van Kan et al., 2017). Despite the comparative efforts reported in **Supplementary Material 2**, the vast majority remain as proteins with hypothetical/unknown biological functions. Therefore, this group of genes represents an opportunity for testing the BEB and determining whether they exhibit particular gene expression profiles shedding light on potential functions.

**Figures 6A and B** show that three genes in each chromosome (Chr17: Bcin17g00150, Bcin17g00160, and Bcin17g00180; Chr18: Bcin18g00090, Bcin18g00120, and Bcin18g00130, respectively) display low mRNA levels across all RNA-Seq libraries. Interestingly, Chr17 genes Bcin17g00010, Bcin17g00020, Bcin17g00040, and Bcin17g00050 display the highest and most likely co-regulated gene expression as deduced by BEB’s clustering function **(Figure 6A)**. As shown in **Supplementary Material 2**, the latter gene encodes a putative NRPS-like protein, whose participation in the biosynthesis of any low molecular weight peptidic product has not yet been described. On the other hand, Chr18 genes Bcin18g00020 and Bcin18g00150 display a distinctive expression pattern during the infection of *A. thaliana* and *S. lycopersicum* **(Figure 6B)**. While these results suggest that these genes might play a role in the infection process, further experimental validation is needed. However, we cannot rule out the possibility that genes with no detectable mRNA levels might be expressed in untested experimental conditions as reported in Aspergillus (Lind et al., 2016).

**Figure 6.**
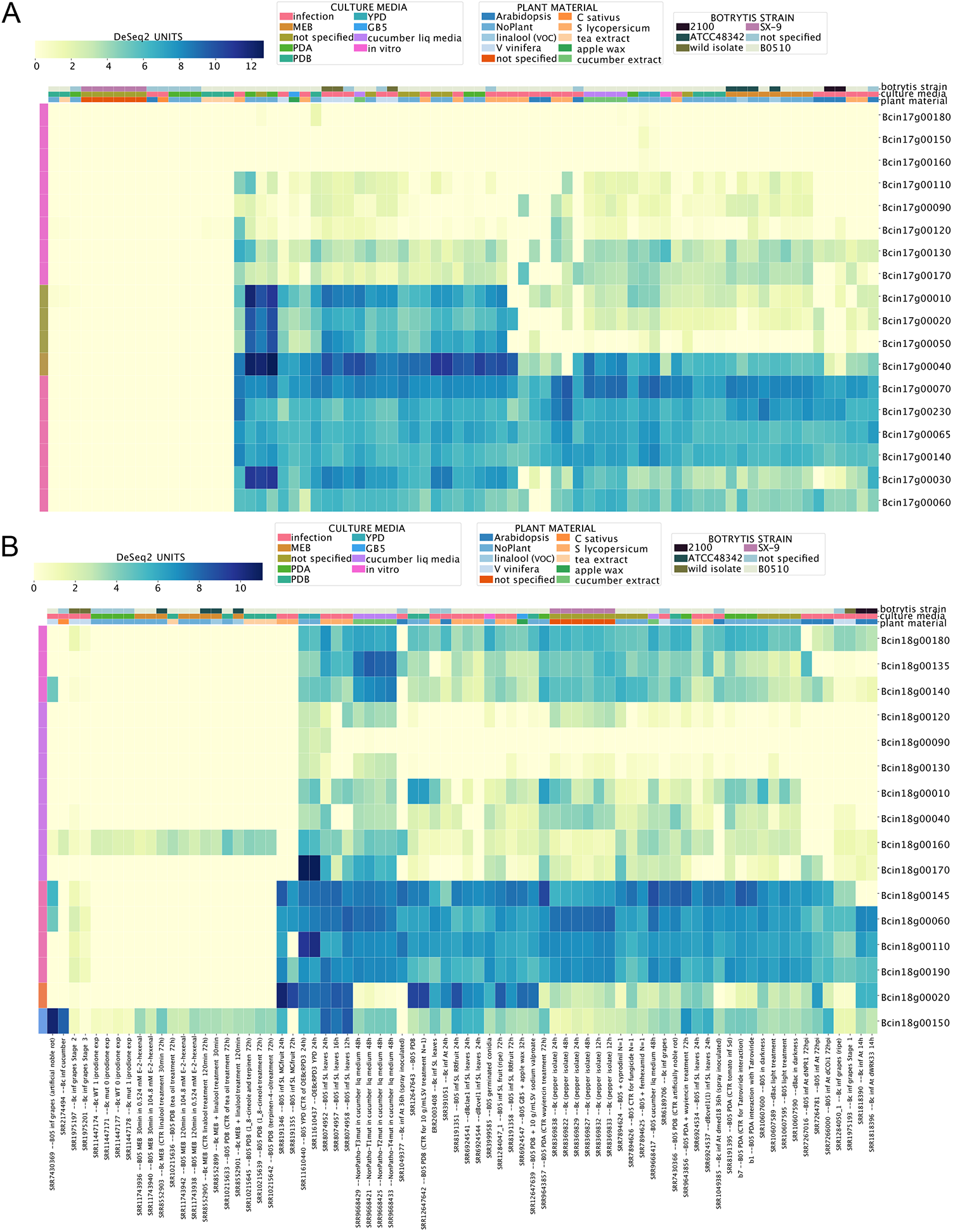
Expression levels of transcripts encoded in mini chromosomes 17 (A) and 18 (B) of *B. cinerea*. Both heatmaps indicate mRNA levels with their corresponding expression scale from yellow to dark blue (low or highly expressed, respectively). The description of each experiment in (A) was omitted to simplify the figure. Both heatmaps indicate mRNA levels and experimental conditions as described in the figure above.

### Inspecting the expression of virulence factors detected by proteomics studies

Considering the relevance of virulence factors in the lifestyle of a phytopathogen such as *B. cinerea*, we next used the BEB to analyze the expression of virulence genes distinct from those required for phytotoxins (described in former sections). In general, virulence factor molecules are extensive in their chemical nature. However, they are characterized by causing harm or suppressing/interfering with host defense strategies, which can also lead to more damage (Pontes et al., 2020). For these reasons, a higher level of expression during the infection of different tissues and plant species is expected. To identify secreted virulence factors, previously published proteomic studies have taken advantage of *in vitro* cultures supplemented with various plant-derived components to induce their expression (Espino et al., 2010; Fernández-Acero et al., 2010; Shah et al., 2009; Shah et al., 2009). The proteins identified from these four publications are summarized in **Supplementary Material 3**. Therefore, we analyzed the expression of their respective genes across experimental conditions using the BEB platform. As depicted in **Supplementary Figure S1 A**, two different expression patterns were observed: i) a high and steady pattern of expression as denoted in the upper section of the heatmap (blue square bracket), and ii) an *in planta* “induced” pattern of expression with two groups of genes indicated with a green and orange square brackets **(Supplementary Figure S1 A)**. In the former group, several unexpected extracellular proteins can be found, including *bcactA* encoding for actin, *bcatp2* coding for the mitochondrial ATP synthase (beta chain), and two additional mitochondrial proteins including malate dehydrogenase and aconitase. Interestingly, *bcspl1* and *bcpg1 —* which play significant roles in virulence *—* were observed in this group (ten et al., 1998; Chagué et al., 2006). Among *in planta* “induced” genes within the green square bracket mentioned above, several glycoside hydrolases were identified (families 5 (three), 6, 7 (two), 10, 11, 28 (two), and 53). There were also two cutinases, and two pectinesterases possibly reflecting a transcriptional regulatory mechanism in this group of carbohydrate-acting enzymes encoding genes **(Supplementary Material 3)**. *In planta* “induced” genes with the orange square bracket **(Supplementary Figure S1 A)** include previously identified virulence factors such as *bcpg3, 4* and *6, bcpgx1, bcxyn11A, bcpme1* and *2*, and *bccutA*, among others. Again, while a common transcriptional regulation seems to be the case as deduced from expression patterns, little is known about the specific TFs controlling the expression of these genes.

### Revisiting the expression of known and proposing new reference genes for transcript level analysis in *B. cinerea*

We integrated and analyzed data spanning a wide variety of experimental conditions **(Supplementary Material 1)**. Thus, we next decided to exploit the wealth of data to revisit the expression of known and previously validated reference genes in *B. cinerea* employed in reverse transcription quantitative real-time PCR (RT-qPCR) studies (Canessa et al., 2013; Ren et al., 2017) and propose new ones. By definition, reference genes tend to be constitutively expressed in all cells of an organism, and their expression levels display slight variations between developmental stages or across various experimental conditions. Therefore, well-normalized RT-qPCR experiments rely on an adequately validated (stable) reference gene(s) (Huggett et al., 2005). In addition, accurate normalization usually requires multiple validated reference genes (Vandesompele et al., 2002), the exact number being dependent on an experimental assessment of variability across the samples of interest (Hellemans and Vandesompele, 2014). Consequently, we took advantage of BEB’s database to determine new reference genes as described in Methods.

We first revisited the expression of seven genes used as a reference in some RT-qPCR assays (Canessa et al., 2013; Ren et al., 2017) **(Table 4** and **Supplementary Material 4)**. See the methods section for details. Bcin11g03430 (*bcsmt3*) and Bcin02g00900 (tubulin, *bctubA*) have a low coefficient of variation (CV) of 0.35 and 0.36, respectively, showing a low disparity of expression levels across all experimental conditions, as expected **(Figure 7)**. In contrast, Bcin15g02120 (CV = 1.4), encoding the commonly-used reference gene *glyceraldehyde-3-phosphate dehydrogenase*, displays obvious changes in expression levels across experiments. Indeed, according to our analysis, it is the worst reference gene of those previously studied **(Table 4** and **Supplementary Material 4)**. In the fungus *Neurospora crassa*, this gene is under the control of its well-known circadian clock (Shinohara et al., 1998), a complex molecular machinery that in *B. cinerea* modulates time-dependent fungal-plant (Hevia et al., 2015)) and fungal-fungal dynamics as recently demonstrated (Henríquez-Urrutia et al., 2022).

**Table 4.**
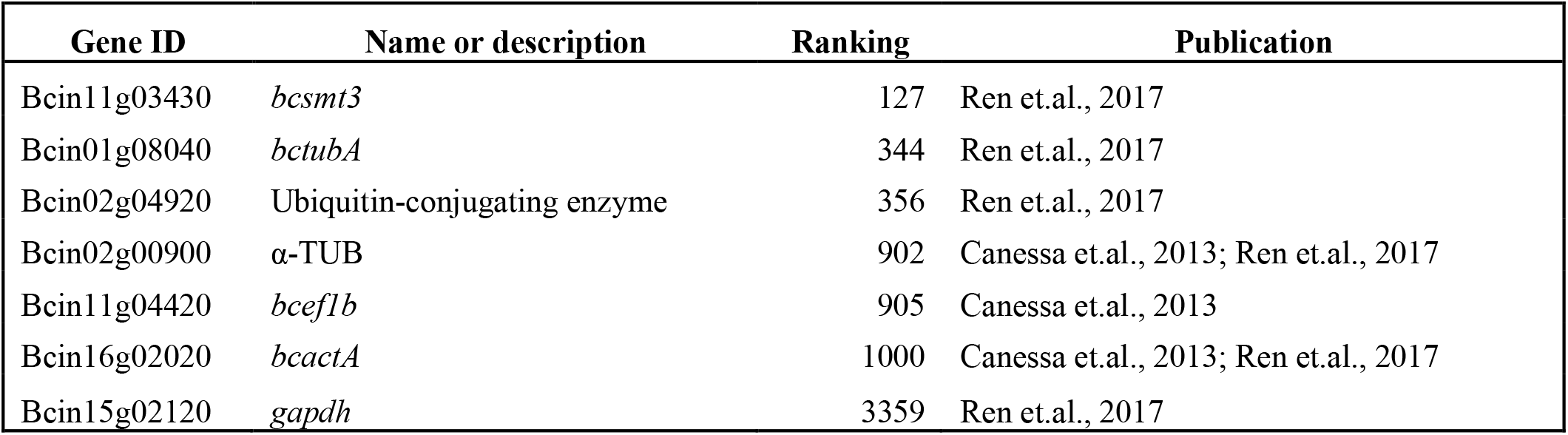
Reference genes validated as such in *B. cinerea* RT-qPCR assays. The rank number was calculated based on the data presented in **Supplementary Material 4**.

**Figure 7.**
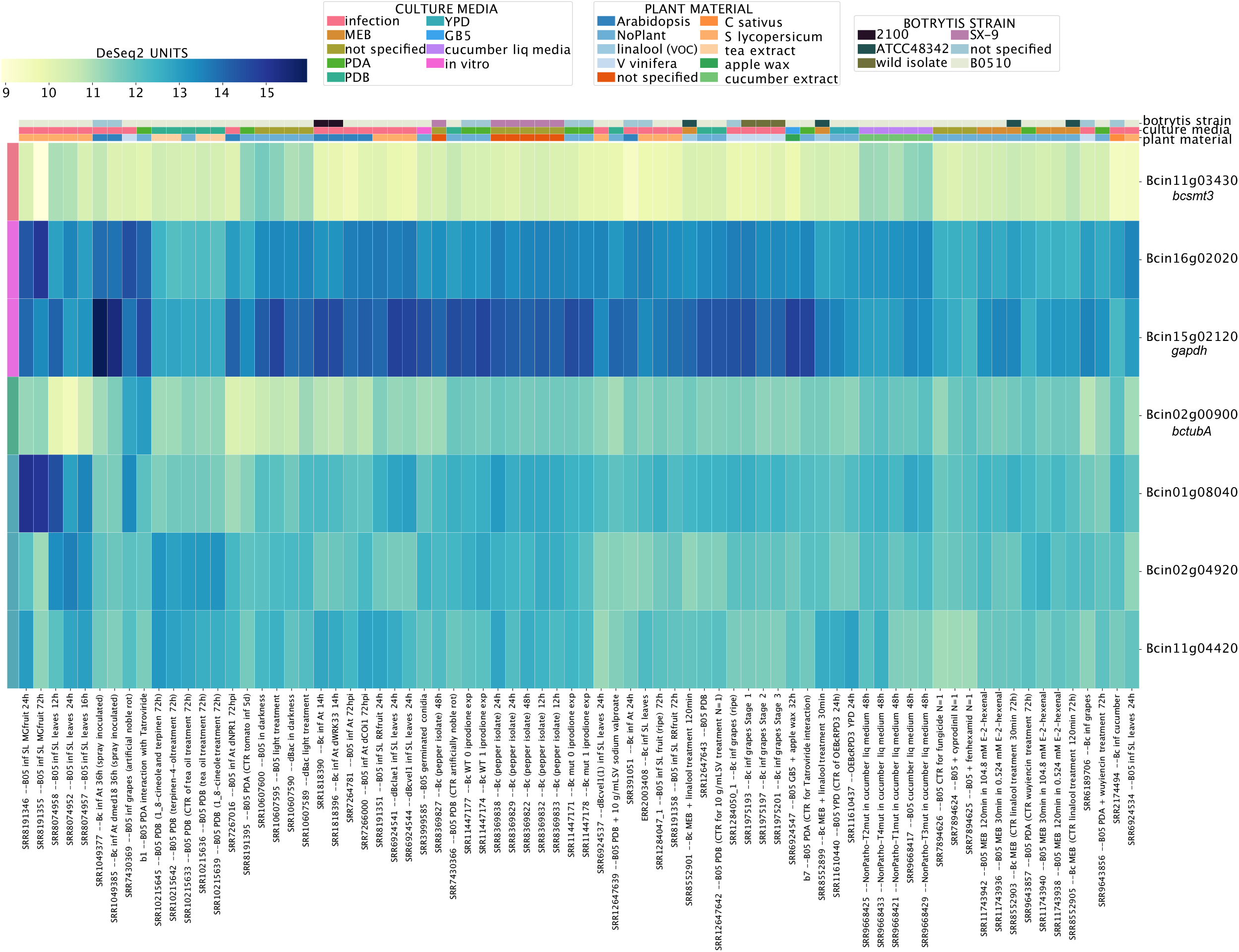
Transcript levels of previously validated reference genes employed in RT-qPCR experiments in *B. cinerea*. The heatmap depicts transcript levels employing the BEB’s continuous color scale from yellow to dark blue (low or highly expressed, respectively). A detailed description of each experiment is provided at the bottom of the heatmap. Experimental conditions and genes are depicted in columns and rows, respectively. Gene IDs are provided at the right of each heatmap.

To provide the *B. cinerea* community with an additional tool, finally, we also generated a list of 40 new reference genes showing the lowest CV across conditions in each global gene expression quartile. Importantly, the proposed reference genes need to be further validated by RT-qPCR, i.e., for an intermediate-low and the intermediate-high expression levels (second and third quartile of normalized expression, respectively) **(Supplementary Material 4)**. Nevertheless, as shown in **Supplementary Figure S2**, their expression is highly similar in different experimental conditions.

## Legend to Supplementary Figures

**Supplementary Figure S1. mRNA levels of virulence factors detected by proteomics studies. (A)** A heatmap depicting the transcript levels of 176 genes employing the BEB’s quartile-categorized expression option (lower: dark green; higher: light green) or the continuous color scale **(B)** from low to high expression (yellow to dark blue). Due to the number of genes being analyzed, neither culture conditions nor gene IDs were not included in the figure, although such information can be found in **Supplementary Table S3**. In (A), genes with a high and steady pattern of expression across conditions are indicated with a blue square bracket, while two groups of genes displaying an *in planta* “induced” pattern of expression are indicated with green and orange square brackets.

**Supplementary Figure S2. mRNA levels of proposed reference genes that can be employed in future RT-qPCR experiments after proper validation**. The heatmap depicts transcript levels employing the BEB’s continuous color scale. Proposed reference genes are indicated in **Supplementary Table 4**. Experimental conditions and gene IDs are indicated in columns and rows, respectively. A steady pattern of expression across conditions is observed for most genes.

## Author Contributions

P.C., D.A., and G.P-L. originated the idea, designed the research, and performed data curation; G.P-L. and P.C. performed data pre-processing and RNA-Seq mapping; D.A., performed software programing; P.C. and G.P-L. wrote original draft including figures with insight from all authors; T.C-M., and J.M-A. performed additional bioinformatic analyses; P.C., G.P-L., D.A., A.V., L.F.L., J.M-A. and R.P. discussed the manuscript and performed final writing, review, and editing; all authors analyzed the experimental data, writing, review and editing.

All authors have read and agreed to the published version of the manuscript.

## Acknowledgments

We sincerely appreciate the leadership, help, and support of our Executive Director at the Millennium Institute for Integrative Biology (iBio), Dra. Susana Cabello.

## Supplementary Material

**Supplementary Material 1**. BEB’s metadata. The table describes the general experimental conditions of each RNA-Seq experiment. Importantly, the table is formatted as on BEB’s server.

**Supplementary Material 2**. Functional annotation of mini-chromosomes (Chr 17 and 18) with protein-encoding genes. The data was obtained after BLAST2GO analysis.

**Supplementary Material 3**. Previously identified extracellular *B. cinerea* proteins detected in *in vitro* cultures supplemented with plant-derived material. The colors of the groups refer to those in Supplementary Figure S1.

**Supplementary Material 4**. Proposed reference genes for RT-qPCR studies. The table shows each gene (ten for each expression quartile), as well as its coefficient of variation (CV).

## Data Availability Statement

The datasets generated for this study can be found in the GitHub (https://github.com/ibioChile/CanessaLab) as well as in the Supplementary Material.

