## Supplementary figures and images for "The *Botrytis cinerea* Gene Expression Browser"

### Supplementary Figure S1

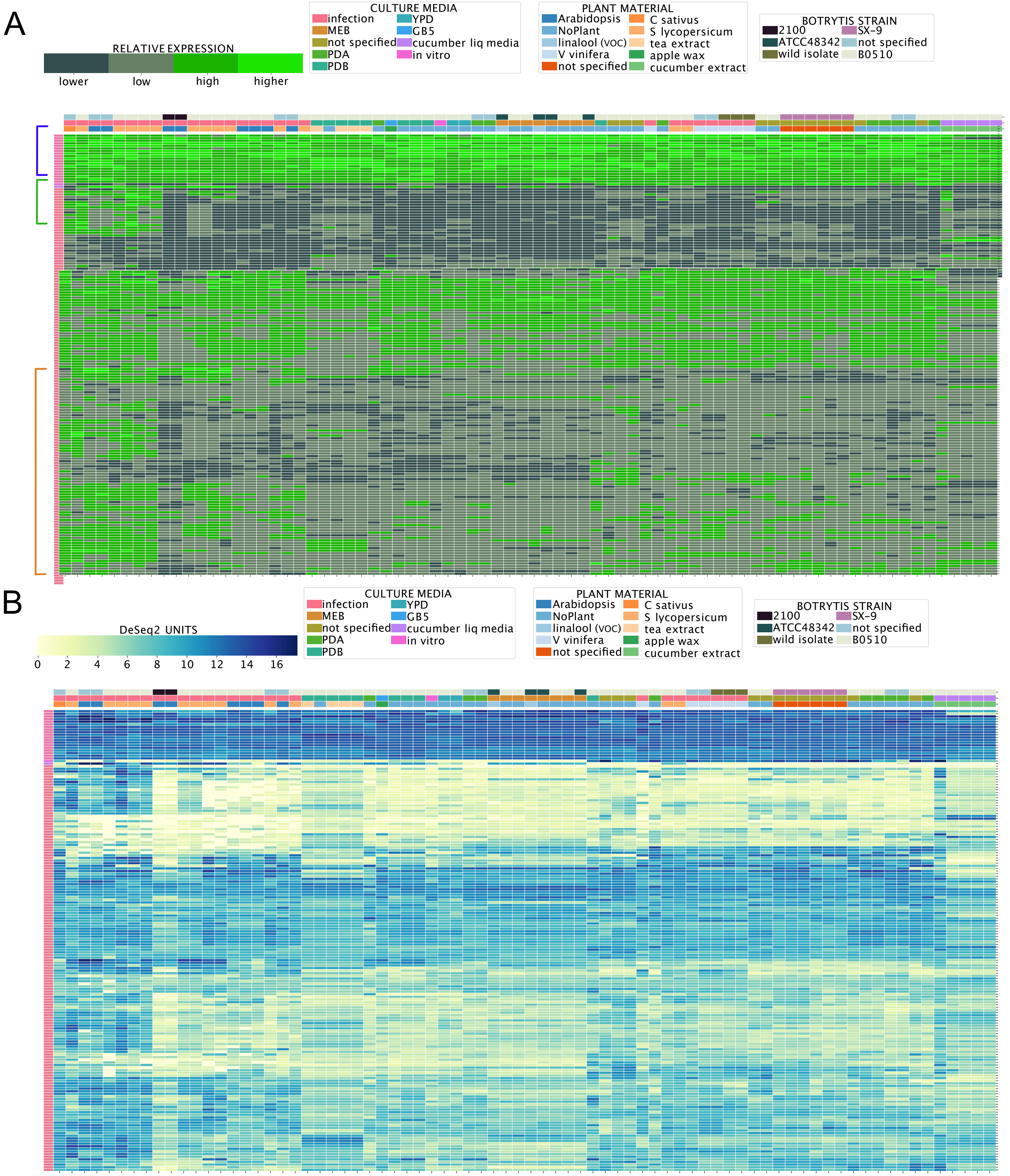

### Supplementary Figure S2

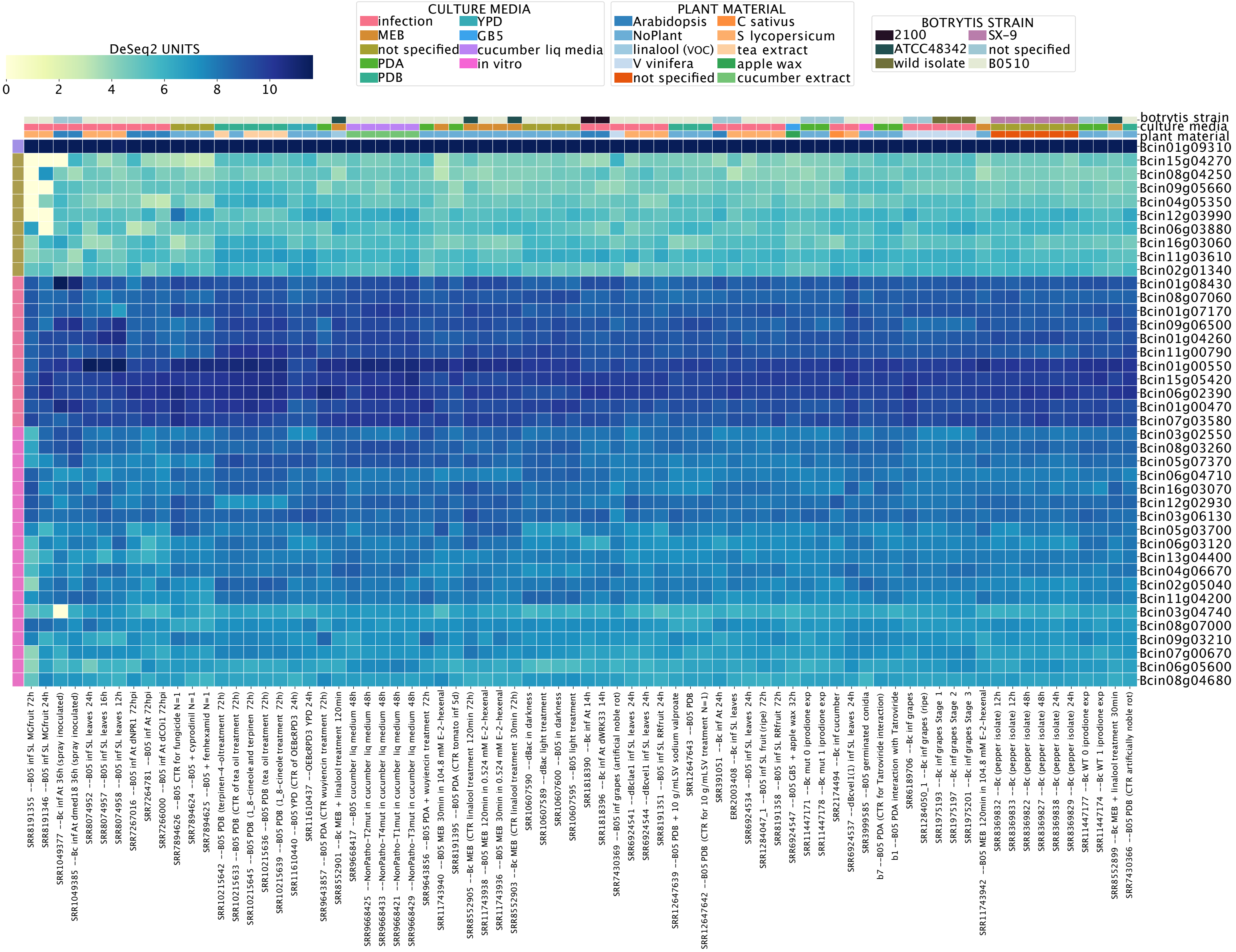
